# Balancing spatial resolution and proteome depth in LC-MS based spatial proteomics

**DOI:** 10.64898/2026.08.27.747491

**Authors:** Mandy Meijer, Jeongwoo Hong, Tobias Pohl, Tomas Koudelka, Claudio Bassot, Hanna Hörnberg, Sumin Lee, Hee Sool Rho, Amos Chungwon Lee, Vicent Pelechano, Ilaria Piazza

## Abstract

Spatial proteomics aims to resolve protein composition within intact tissues, yet extraction-based liquid chromatography–mass spectrometry (LC–MS) workflows face an inherent trade-off: smaller sampling units increase spatial specificity, whereas larger sampling units provide greater proteome depth and robustness. As analytical sensitivity improves, sampling-unit size therefore becomes a key experimental design parameter.

Current extraction-based LC-MS workflows typically rely on laser capture microdissection (LCM), where sample recovery and scalability can become limiting at low input. Spatially resolved laser-activated cell sorting (SLACS) offers an alternative tissue-isolation strategy based on single-pulse near-infrared laser activation. Here, we use SLACS to systematically examine the resolution– sensitivity trade-off across sampling units ranging from single-cell-equivalent to larger low-input tissue regions. Few-cell sampling retained substantial proteomic information relative to larger regions while increasing spatial specificity. Applied to the mouse somatosensory cortex, SLACS generated deep, layer-resolved proteomic profiles from regions corresponding to approximately 60 cells and preserved major layer-specific molecular patterns at inputs as low as approximately 6 cells. These results highlight sampling-unit size as an important experimental design parameter in extraction-based spatial proteomics and support few-cell sampling as a practical compromise between spatial specificity, proteome depth and robustness.

## Main

Spatial organization is central to cellular identity and function, driving the need for molecular measurements within intact tissue contexts. While spatial transcriptomics has enabled large-scale mapping of gene expression patterns, proteins constitute a major functional layer of cellular regulation and are regulated beyond RNA abundance. Accordingly, spatial proteomics provides a more direct molecular readout of cellular state. However, unlike nucleic acids, proteins cannot be enzymatically amplified, imposing stringent requirements on sample integrity, isolation efficiency, and analytical sensitivity, particularly as the sampled tissue regions become smaller.

Current spatial proteomics approaches broadly fall into two categories. Imaging-based approaches measure proteins directly within intact tissue sections and include affinity-based imaging methods, ranging from conventional immunofluorescence and immunohistochemistry to highly multiplexed platforms (Uhlen *et al*., 2005; Uhlen *et al*., 2015; Angelo *et al*., 2014; Giesen *et al*., 2014; Goltsev *et al*., 2018; Lin *et al*., 2018), as well as mass spectrometry imaging approaches such as MALDI-MSI (Lopez & Hummon, 2026). These methods preserve continuous spatial information but differ in molecular coverage, sensitivity and degree of targeting. A second category comprises extraction-based approaches, in which defined tissue regions are isolated for downstream liquid chromatography-mass spectrometry (LC-MS) analysis (Zhu *et al*., 2018; Xu *et al*., 2018; Mund *et al*., 2022). These workflows have evolved from early low-input approaches, such as nanoPOTS (Zhu *et al*., 2018) and SIS-PROT (Xu *et al*., 2018), to more recent developments such as deep visual proteomics (DVP), which combines microscopy-guided tissue selection with image analysis and deep LC-MS-based proteome profiling (Mund *et al*., 2022). Extraction-based approaches enable deep and relatively unbiased proteome profiling but physically remove sampled regions from their tissue context, thereby disrupting the continuous spatial organization of the tissue. Spatial relationships must therefore be reconstructed from independently measured regions rather than directly observed. The size of the isolated tissue region therefore becomes a central experimental parameter: smaller sampling units increase spatial specificity but reduce the amount of material available for LC-MS analysis, whereas larger sampling units improve proteome depth and reproducibility but average across more cells and reduce spatial resolution.

Recent advances in LC-MS sensitivity and throughput now enable deep proteome profiling from very small amounts of material, down to tens of cells or fewer (Budnik *et al*., 2018; Brunner *et al*., 2022; Leduc *et al*., 2022), including spatially resolved single cells (Rosenberger *et al*., 2023). As analytical sensitivity improves, however, the central question shifts from whether such small samples can be measured to how small a tissue region should be sampled to obtain biologically meaningful and reproducible information. Single-cell analysis provides maximal spatial specificity, but individual cells do not necessarily constitute equivalent proteomic sampling units: cell types vary widely in size, morphology, protein content, and contribution from surrounding cellular structures, particularly in heterogeneous tissues such as the brain. Thus, for many spatial proteomics applications, the most informative sampling unit may not be a single cell, but rather a small tissue area that balances spatial specificity with sufficient material while reducing variability in total protein input.

In extraction-based spatial proteomics, the practical performance of small sampling units also depends on how reliably the selected tissue can be isolated and recovered. Laser capture microdissection (LCM), in which selected tissue regions are typically excised from polymer membrane slides using an ultraviolet (UV) laser, is widely used for this purpose, but incomplete detachment or recovery of excised tissue fragments can become increasingly problematic as sampling-unit size decreases. At low input, failure to recover a selected region may result in loss of the entire sample, making collection efficiency an important determinant of workflow robustness, particularly for rare or spatially restricted tissue regions.

Spatially resolved laser-activated cell sorting (SLACS) provides an alternative tissue isolation strategy based on near-infrared (near-IR) laser activation rather than perimeter cutting (Jeong et al., 2023). Tissue sections are mounted on membrane-free, metal oxide-coated slides, and selected regions are released in a single laser pulse directly into collection wells. SLACS has previously been applied to genomic, transcriptomic and epitranscriptomic analyses (Jeong et al., 2023; Lee et al., 2022). Its membrane-free, single-pulse distinct release mechanism therefore provides a useful framework for testing whether robust tissue recovery can be maintained as sampling-unit size decreases.

Here, we use SLACS to examine sampling-unit size as an explicit experimental design parameter in extraction-based spatial proteomics. Rather than treating single-cell profiling as the endpoint, we systematically evaluate how decreasing sampling-unit size affects proteome depth, reproducibility, collection robustness, and preservation of biological information. We first benchmark sampling units spanning single-cell, few-cell, and larger low-input regions, using comparison with UV-LCM to establish the collection robustness of SLACS. We then apply SLACS to the mouse somatosensory cortex, a structurally and molecularly complex tissue with well-defined cortical layers, to test whether biologically meaningful spatial proteomic information is preserved as sampling-unit size is reduced. By comparing larger cortical regions with few-cell regions, we show that few-cell sampling can provide a useful compromise between spatial specificity, proteome depth and robustness while preserving layer-specific proteomic organization.

## Results

### Sampling-unit size defines a resolution-sensitivity trade-off in deep spatial proteomics

Deep spatial proteomics requires choosing the size of the tissue region to be isolated for LC-MS analysis. This sampling-unit size determines the balance between spatial specificity and the amount of proteomic material available for measurement. Single-cell analysis provides maximal spatial resolution but is technically demanding and highly sensitive to differences in cell size, morphology, and protein content. Importantly, different cell types contribute vastly different amounts of protein, resulting in large differences in the total proteomic material entering the analysis. Because mass spectrometry relies on analyzing comparable amounts of material, using single cells as input introduces a fundamental challenge: individual cells may not represent equivalent analytical sampling units, complicating quantitative comparison and data normalization. If the primary goal is to resolve spatial organization, pushing the sampling unit down to individual cells may therefore become a complication rather than an advantage. Conversely, larger areas improve proteome depth and reproducibility but average across more cells, reducing the ability to resolve fine spatial heterogeneity. We therefore reasoned that sampling units corresponding to a few cells may provide a practical compromise between spatial specificity and a more comparable proteomic input **(Fig. 1a).**

**Figure 1.**
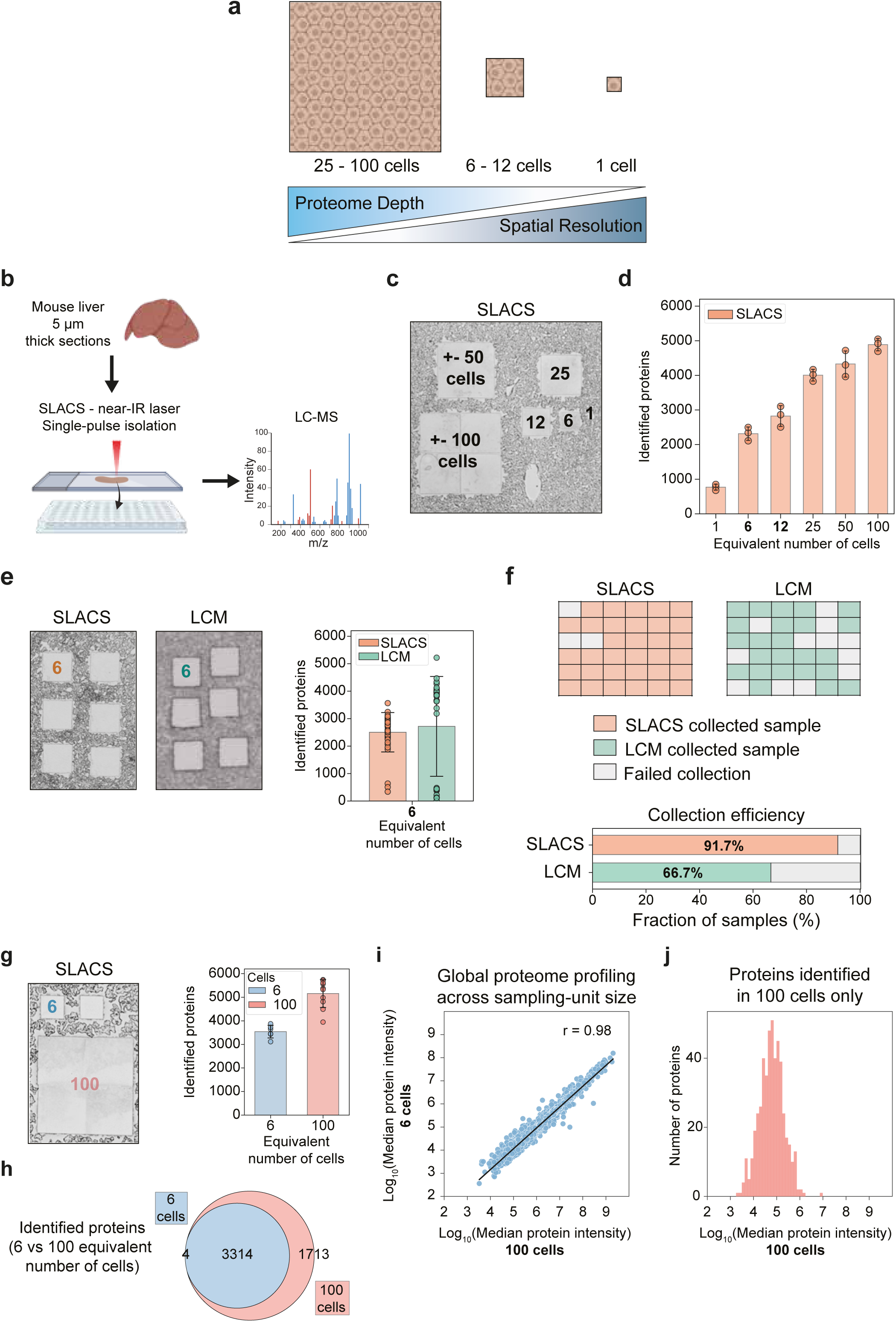
**Benchmarking sampling unit-size for LC-MS-based spatial proteomics**. **a**, Conceptual overview of sampling-unit sizes in LC-MS-based spatial proteomics, ranging from low-input areas (25 - 100 cells equivalent), to few-cell (6 - 12 cells equivalent) and single-cell (1 cell equivalent) sampling units, illustrating the trade-off among proteome depth, spatial specificity, and sampling robustness. **b**, Workflow for isolation of tissue areas from mouse liver using spatially resolved laser-activated cell sorting (SLACS). Tissue regions are released by a single near-IR laser pulse, collected into 96-well plates, and processed by in-well lysis and digestion prior to DIA-MS analysis to assess proteome coverage across different sampling-unit sizes. *Created with Biorender.com*. **c**, SLACS isolation of liver tissue areas of 600 μm^2^, 6,000 μm^2^, 12,500 μm^2^, 25,000 μm^2^, 50,000 μm^2^ and 100,000 μm^2^ equivalent to approximately 1, 6, 12, 25, 50, and 100 cells. Wide-field image of the liver tissue after isolation. **d**, Number of identified protein groups per collected input size using SLACS. Bars represent the mean and error bars indicate SD (n=3). **e**, Representative images of liver tissue after isolation of 6,000 μm^2^ areas (equivalent to approximately 6 cells) using SLACS and UV-LCM (left), and the corresponding protein group identifications across 36 replicates (right). Bars represent the mean and error bars indicate SD. **f**, Collection efficiency of SLACS and UV-LCM for 36 replicate isolations of 6,000 μm^2^ areas (equivalent to approximately 6 cells). **g**, Representative images of liver tissue after isolation of 6,000 μm^2^ and 100,000 μm^2^ areas (equivalent to approximately 6 and 100 cells, respectively) using SLACS (left), and the corresponding protein group identifications (right). Data represent 6 replicates for the 6-cell sampling unit and 10 replicates for the 100-cell sampling unit. Bars represent the mean and error bars indicate SD. **h,** Overlap of proteins identified in the 6- and 100-cell SLACS datasets in panel g. Data represent 6 - 10 replicates per collected input size. **i**, Protein abundance is preserved across sampling unit size: Scatter plot comparing median protein intensities measured with DIA-MS between 100-cell and 6-cell SLACS samples. Each point represents one protein quantified at both sampling-unit sizes using the median log_10_-transformed intensity across 6 replicates for the 6-cell sampling unit and 10 replicates for the 100-cell sampling unit. Protein abundances were highly correlated across sampling-unit sizes (Correlation coefficient r = 0.98; Pearson correlation). **j,** Distribution of protein abundances for proteins identified in 100-cell SLACS samples but not in 6-cell samples. Intensities correspond to the median protein abundance in the 100-cell dataset. Proteins absent at the 6-cell sampling-unit size were predominantly of low abundance, indicating that reducing sampling-unit size primarily affects detection sensitivity.

To evaluate this trade-off systematically, we analyzed defined tissue regions spanning single-cell, few-cell, and larger low-input sampling units using SLACS coupled to LC-MS (**Fig 1a and 1b**). Using mouse liver tissue as a relatively homogeneous benchmarking system (Makhmut *et al*., 2023), we isolated regions with an area estimated to contain 1 cell (600 µm²), 6 cells (6,000 µm²), 12 cells (12,500 µm²), 25 cells (25,000 µm²), 50 cells (50,000 µm²), and 100 cells (100,000 µm²) (**Fig 1c**). These sampling units were selected to cover the transition from technically challenging single-cell input to larger regions that provide higher sensitivity but reduced spatial specificity.

Across input sizes, the number of identified protein groups increased with sampling-unit size, as expected from the greater amount of material available for LC-MS analysis (**Fig 1d**). The mean number of identified protein groups increased from 770 for single-cell sampling units to 4,887 for 100-cell sampling units (n = 3 per group) (**Fig 1d**). The few-cell range between 6 and 12 cells retained substantial proteome coverage within a range between 2,317 to 2,825 proteins (**Fig 1d**), suggesting that sampling units corresponding to several cells may provide a practical compromise between spatial specificity and analytical sensitivity. These observations motivated a more detailed assessment of collection robustness and proteomic reproducibility in this few-cell regime.

### SLACS improves recovery of few-cell sampling units

Because robust isolation becomes increasingly important as sampling-unit size decreases, we evaluated SLACS as a tissue collection strategy for few-cell spatial proteomics. In contrast to UV- LCM, where tissue areas are isolated by perimeter cutting from polymer membrane slides, SLACS uses single-pulse near-IR laser activation to release tissue regions from membrane-free, metal oxide-coated slides directly into collection wells (**Extended Data Fig 1a and 1b**). This difference in tissue-release mechanism is expected to reduce sample loss and improve collection robustness, particularly for small sampling units.

At few-cell input, incomplete tissue recovery can dominate measurement quality because failed collections result in the loss of entire samples. To directly assess this limitation, we compared collection efficiency between SLACS and UV-LCM using replicate areas corresponding to approximately 6 cells. This sampling-unit size was chosen because it represents a regime in which spatial specificity is substantially increased relative to larger regions, while still providing more proteomic material than single-cell sampling (**Fig 1d**). UV-LCM and SLACS yielded broadly comparable proteome profiles, with SLACS identifying 2,506 ± 717 and UV-LCM 2,719 ± 1,817 protein groups (mean ± s.d.; n = 36 per method) (**Fig 1e**). Although UV-LCM yielded slightly higher protein group identification numbers in successfully collected samples, it also exhibited substantially greater variability, indicative of less consistent tissue collection (**Fig 1e**). Among 36 replicate 6-cell areas, 12 UV-LCM collections failed, compared to only 3 SLACS collections, corresponding to collection efficiencies of 66.7% and 91.7%, respectively (**Fig 1f**). Thus, although UV-LCM can provide high proteome depth when collection succeeds, SLACS provides more reliable recovery of few-cell sampling units. Additional experiments with both Orbitrap Astral and TimsTOF SCP mass spectrometers yielded similar results (**Extended Data Fig 1c and 1d**), suggesting that the observed effects are largely independent of MS performance and instead reflect differences in the sampling isolation and collection workflow.

These results indicate that workflow performance at small sampling-unit size cannot be judged solely by protein identifications in successfully measured samples. Instead, collection efficiency and dropout become central workflow metrics. By reducing sample loss, SLACS enables more reproducible operation in the few-cell regime and provides a practical strategy for sampling small or spatially restricted tissue regions.

### Few-cell spatial sampling preserves proteome profiles relative to larger sampling units

Having established improved collection robustness with SLACS, we next asked how reducing sampling-unit size affects proteome coverage and the preservation of quantitative information. For spatial proteomics, the key question is not only how many proteins can be identified, but how much of the proteomic information obtained from larger regions is retained at higher spatial resolution. We therefore compared SLACS-isolated 100-cell and 6-cell areas. Despite an approximately 17- fold reduction in sampled area, 3,314 of the 5,027 protein groups consistently detected in 100-cell samples were also identified at 6-cell input (**Fig. 1g and h**), demonstrating that a substantial fraction of the proteome is retained in few-cell sampling units.

Protein abundances remained highly correlated between 100-cell and 6-cell areas (Pearson correlation, r = 0.98; **Fig. 1i**), although overall signal intensity decreased as expected at lower input. Proteins detected in 100-cell samples but not at 6-cell input were predominantly of low abundance, indicating that reducing sampling-unit size primarily limits detection sensitivity rather than broadly altering the measured proteome (**Fig. 1j**). Reproducibility remained high within both sampling-unit sizes (**Extended Data Fig. 1e**). Together, these results indicate that few-cell sampling preserves much of the quantitative proteomic information obtained from larger regions while providing substantially greater spatial specificity.

### Few-cell sampling preserves cortical layer organization

We next asked whether the proteomic information retained at few-cell input is sufficient to resolve biologically meaningful spatial organization in a complex tissue by comparing the expected relative abundance profiles of known protein markers. The mouse somatosensory cortex provides a stringent test system because cortical layers differ in cellular composition, connectivity, and function across relatively small spatial distances. We therefore used SLACS to isolate regions from layers 2/3, 4, 5, and 6, performed a differential protein abundance analysis using DIA-MS as readout, and compared results from large and few-cell sampling unit sizes (**Fig 2a**).

**Figure 2.**
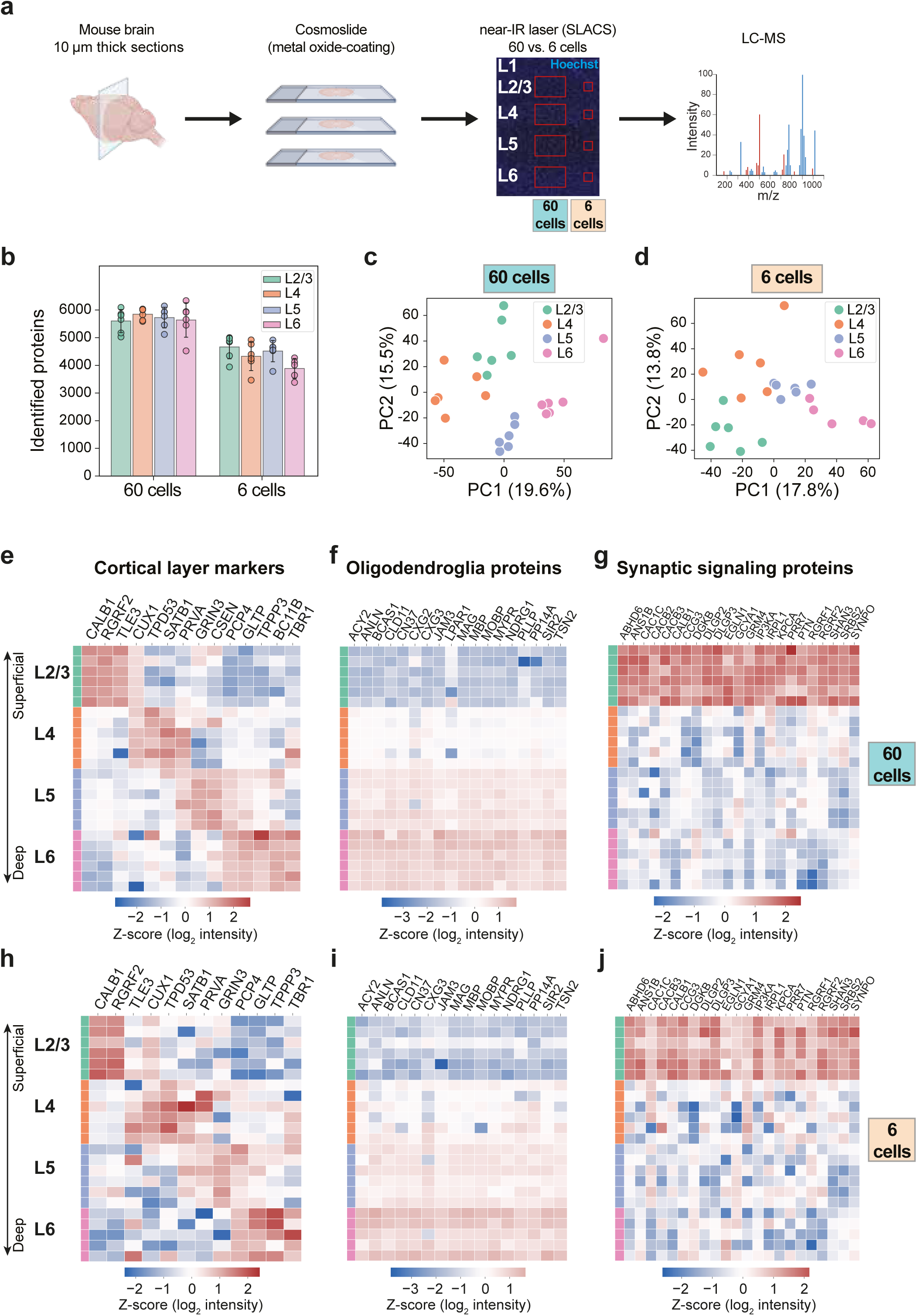
Few-cell sampling preserves cortical layer organization. **a**, Schematic workflow for isolation of 200 x 300 μm (equivalent to approximately 60 cells) and 77 x 77 μm (equivalent to approximately 6 cells) areas from the mouse somatosensory cortex for spatial proteomic analysis. Mouse brains are cryosectioned into 10 μm-thick sections, and cortical layer areas are isolated from adjacent sections using near-IR laser pulses. Isolated areas are collected into 96-well plates for in-well lysis and digestion prior to DIA-MS analysis, label-free quantification, and pairwise differential abundance analysis between cortical layers. *Created with Biorender.com*. **b**, Number of identified protein groups per cortical layer for the 60-cell and 6-cell sampling-unit sizes. Data represent 5-6 replicates per cortical layer. Bars represent the mean and error bars indicate SD. **c**, Principal component analysis (PCA) of scaled protein abundances across cortical-layer samples at the 60-cell sampling unit size. Each point represents one replicate. **d**, PCA of scaled protein abundances across cortical-layer samples at the 6-cell sampling unit size. Each point represents one replicate. **e**, Heatmap of annotated cortical layer marker proteins showing significant differential abundance in at least one pairwise comparison between cortical layers (L2/3, L4, L5 and L6) in the 60-cell dataset (adjusted p < 0.05, Benjamini-Hochberg FDR correction, |log_2_ fold change| > 0.58). Columns represent cortical layer marker proteins, and rows represent individual cortical layer replicates of 60 cell equivalents (n = 6). Relative protein abundances are shown as z-score-scaled intensities for each protein group across all samples. **f**, Heatmap of annotated oligodendroglia proteins significantly enriched in layer 6 relative to layer 2/3 in the 60-cell dataset (adjusted p < 0.05, Benjamini-Hochberg FDR correction, |log_2_ fold change| > 0.58). Columns represent oligodendroglia proteins, and rows represent individual cortical layer replicates of 60 cell equivalents (n = 6). Relative protein abundances are shown as z-score-scaled intensities for each protein group across all samples. **g**, Heatmap of proteins associated with synaptic signaling showing significantly higher abundance in layer 2/3 than in each of the other cortical layers (L4, L5, and L6) in the 60-cell dataset (adjusted p < 0.05, Benjamini-Hochberg FDR correction, |log_2_ fold change| > 0.58). Columns represent synaptic signaling proteins, and rows represent individual cortical layer replicates of 60 cell equivalents (n = 6). Relative protein abundances are shown as z-score-scaled intensities for each protein group across all samples. **h**, Heatmap of annotated cortical layer marker proteins as shown in Fig. 2e, displayed for the 6- cell dataset. Columns represent cortical layer marker proteins, and rows represent individual cortical layer replicates of 6 cell equivalents (n = 5 or 6). Relative protein abundances are shown as z-score-scaled intensities for each protein group across all samples. **i**, Heatmap of annotated oligodendroglia proteins as shown in Fig. 2f, displayed for the 6-cell dataset. Columns represent oligodendroglia proteins, and rows represent individual cortical layer replicates of 6 cell equivalents (n = 5 or 6). Relative protein abundances are shown as z-score- scaled intensities for each protein group across all samples. **j**, Heatmap of proteins associated with synaptic signaling as shown in Fig. 2g, displayed for the 6- cell dataset. Columns represent synaptic signaling proteins, and rows represent individual cortical layer replicates of 6 cell equivalents (n = 5 or 6). Relative protein abundances are shown as z- score-scaled intensities for each protein group across all samples.

We first analyzed larger cortical sampling units of 60,000 µm², corresponding to approximately 60 cells per area (**Extended Data Fig 2a**). SLACS enabled deep and consistent proteome coverage, with an average of 5,703 protein groups identified per sample (**Fig 2b**). Principal component analysis (PCA) revealed clear separation of cortical layers (**Fig 2c**), with most replicates clustering by layer, indicating that layer-specific proteomic signatures are captured reproducibly. An independent replicate experiment yielded comparable proteome depth and high overlap between datasets (**Extended Data Fig 2b-d**), further supporting reproducibility of the workflow.

To test whether this spatial proteomic information is preserved at higher spatial sampling resolution, we then analyzed substantially smaller cortical areas of 6,000 µm², corresponding to approximately 6 cells (**Extended Data Fig 2a**). Despite the reduced sampling-unit size, SLACS maintained robust proteome coverage, with an average of 4,367 protein groups identified per sample (**Fig 2b**). Only one of 24 collected regions was classified as a failed collection, corresponding to a dropout rate of 4.2%. PCA analysis revealed a layer-dependent gradient, indicating that cortical layer structure remains detectable at few-cell input (**Fig 2d**).

Direct comparison of the two sampling-unit sizes showed that 6-cell areas shared approximately 80% of identified proteins with 60-cell regions (4,667 over 5,798 proteins **Extended Data Fig 2e**). Layer-resolved differences were also limitedly preserved, with 557 significantly changing proteins overlapping between datasets (**Extended Data Fig 2e**). These results show that reducing the sampling unit from approximately 60 to 6 cells preserves a substantial fraction of both proteome coverage and information about the spatially structured protein variation in the somatosensory cortex.

### Spatial proteomic signatures are retained at few-cell resolution

To determine whether preserved layer separation reflected meaningful biological organization, we examined known cortical layer markers and spatially patterned protein groups across the two sampling-unit sizes. In the 60-cell dataset, differential protein abundance analysis recapitulated established cortical layer markers, including RGRF2 (RASGRF2) and CALB1 in layers 2/3, CUX1 in layers 2-4, PRVA (PVALB) in layers 4/5, BC11B (Bcl11b/CTIP2) in layers 5/6, and TBR1 in layer 6 (Alcantara *et al*., 1993; Bulfone *et al*., 1995; Leid *et al*., 2004; Nieto *et al*., 2004; Zeng *et al*., 2012) (**Fig 2e; Supplementary Table 1**). Additionally, proteins associated with oligodendroglia and myelination showed a gradual increase from superficial to deep layers (**Fig 2f**), consistent with the higher density of myelinated axons in deeper cortical regions (Parnavelas *et al*., 1983).

We next asked whether layer-specific protein differences reflected coordinated functional programs rather than isolated marker proteins. Gene Ontology (GO) enrichment analysis identified synaptic signaling as a major functional category among significantly differential proteins (**Extended Data Fig 2f**). Consistent with the prominent local circuitry of superficial cortical layers, 12.5% of differentially abundant synaptic proteins were specifically enriched in layers 2/3 (**Fig 2g**). More broadly, synaptic proteins, including neurotransmitter receptors, ion channels, and signaling proteins displayed distinct layer-dependent expression profiles across cortical layers (**Supplementary Table 1**). Collectively, these results demonstrate that SLACS captures layer-specific synaptic specialization.

Importantly, these spatial signatures were retained in the 6-cell dataset. Most canonical layer markers were robustly detected and showed the same layer-dependent patterns observed in larger regions (**Fig 2h, Supplementary Table 2**). Oligodendroglia and myelin-associated proteins similarly increased toward deeper layers (**Fig 2i**), and proteins associated with synaptic signalling in layer 2/3 in the 60-cell dataset largely retained enrichment in superficial layers (**Fig 2j**). Thus, few-cell sampling preserves not only global layer separation but also specific protein programs that reflect known cortical organization.

These findings support the view that few-cell sampling units provide sufficient proteomic depth to recover meaningful spatial biology while increasing spatial specificity relative to larger regions. The preservation of cortical layer markers, myelin-associated gradients, and synaptic protein patterns indicates that the few-cell regime captures biologically interpretable spatial proteomic information rather than merely producing reduced-depth measurements.

### Reducing sampling-unit size reveals the sensitivity cost of higher spatial resolution

Although few-cell sampling preserved major spatial proteomic features, reducing sampling-unit size also imposed measurable costs. The number of quantified protein groups decreased from an average of 5,703 in 60-cell cortical areas to 4,367 in 6-cell areas (**Fig 2b**). Only 557 over 1,320 (for 60 cells) and over 847 (for 6 cells) differentially abundant proteins were identified by both 60 and 6 cells datasets (**Extended Data Fig 2e**). Proteins lost at smaller sampling-unit size were enriched among lower-abundance proteins (**Extended Data Fig 2g and 2h**), consistent with detection sensitivity becoming limiting as input decreases (**Fig 1j**). In addition, although median coefficients of variation were broadly comparable between 60-cell and 6-cell areas, some layers showed increased variability at lower input (**Extended Data Fig 2i**).

Functional enrichment analysis showed that differentially abundant proteins in both the 60-cell and 6-cell datasets were enriched for synaptic signaling and other key layer-associated functional programs (**Extended Data Fig 2j**). In contrast, proteins showing significant differential abundance only in the 60-cell dataset were associated with broader cellular processes (**Extended Data Fig 2j**), suggesting that reduced sampling-unit size preferentially affects protein sensitivity, while preserving biological information about spatially organized biological programs.

K-means clustering of significantly differential proteins further supported this interpretation. Both 60-cell and 6-cell datasets revealed five layer-dependent expression signatures (**Fig 3a and 3b**), including synaptic programs, layer 4-enriched metabolic and chromatin-associated signatures, and a deep-layer myelin/neurofilament-associated program (**Extended Data Fig 3a and 3b**). Both the expression patterns and underlying biological programs showed strong overlap. Thus, reducing sampling-unit size preserves the dominant spatially organized proteomic programs while reducing detection of lower-abundance proteins and less robust features.

**Figure 3.**
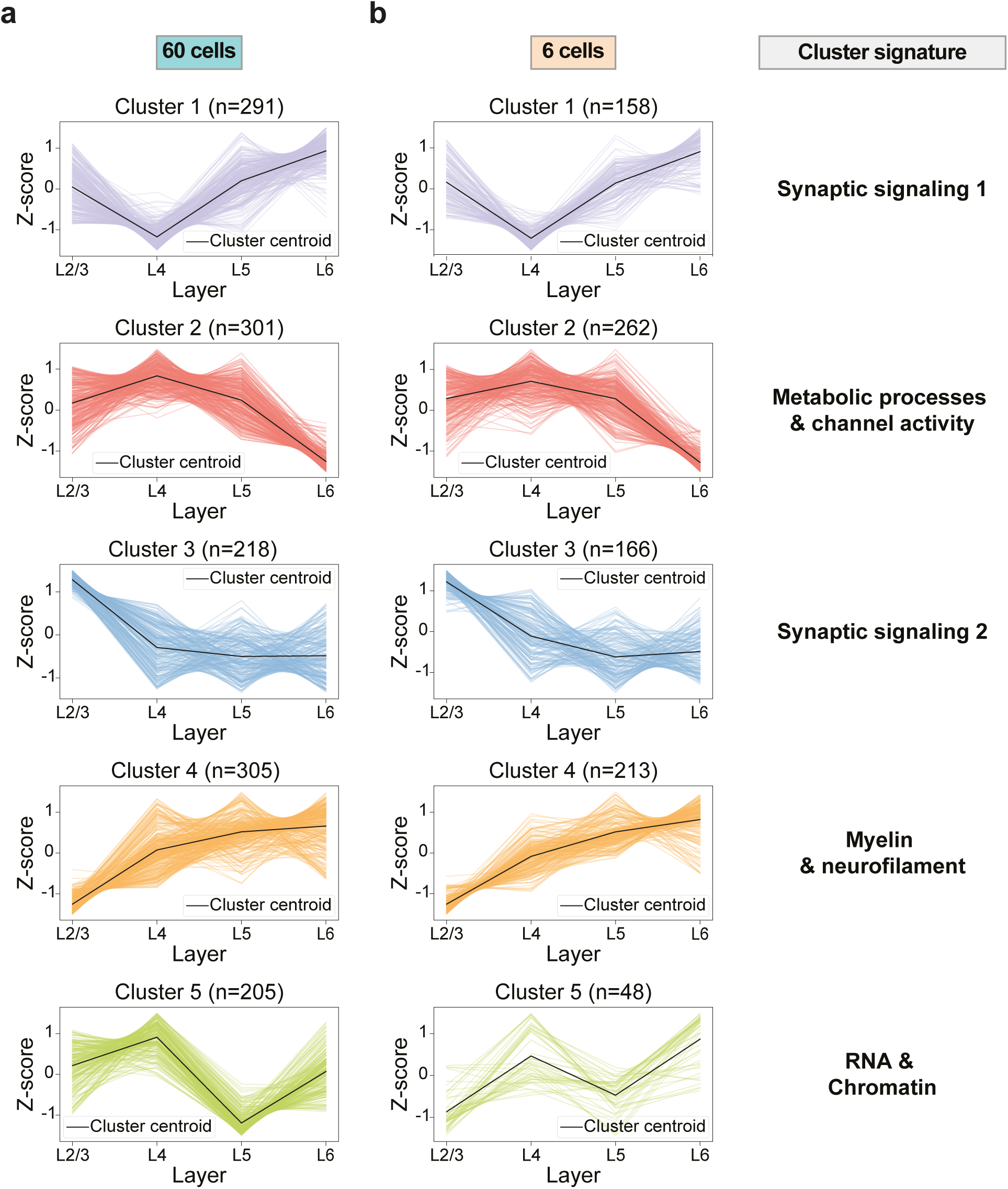
Layer-specific proteomic signatures at few-cell input. **a**, K-means clustering of proteins showing significant differential abundance in at least one pairwise comparison between cortical layers (n = 1,320) in the 60-cell dataset, identified using linear models with empirical Bayes moderation (adjusted p < 0.05, Benjamini-Hochberg FDR correction, |log_2_ fold change| > 0.58), based on row-wise z-score-scaled protein abundances across cortical layers. Five distinct layer-dependent expression signatures were identified. Colored lines represent individual proteins, and black lines indicate cluster centroids (mean profiles). **b**, K-means clustering of proteins showing significant differential abundance in at least one pairwise comparison between cortical layers (n = 847) in the 6-cell dataset, identified using linear models with empirical Bayes moderation (adjusted p < 0.05, Benjamini-Hochberg FDR correction, |log_2_ fold change| > 0.58), based on row-wise z-score-scaled protein abundances across cortical layers. Five distinct layer-dependent expression signatures were identified. Colored lines represent individual proteins, and black lines indicate cluster centroids (mean profiles).

Together, these analyses quantify the trade-off associated with increasing spatial resolution. Few-cell sampling units provide substantially higher spatial specificity than larger regions while retaining reproducible detection of major biological programs. At the same time, they incur a sensitivity cost, particularly for low-abundance proteins. In the systems tested here, few-cell sampling therefore provides a useful compromise between spatial specificity, proteome depth and robustness, particularly for rare or spatially restricted tissue regions.

## Discussion

A central challenge in extraction-based spatial proteomics is choosing the size of the tissue area to analyze. Smaller sampling units increase spatial specificity but reduce the amount of material available for LC-MS analysis, whereas larger sampling units improve proteome depth and reproducibility but average over more cells. Here, we systematically evaluated this resolution/sensitivity trade-off and show that few-cell sampling units can provide a practical operating regime for deep spatial proteomics. Regions corresponding to approximately 6 cells retained substantial proteome coverage, reproducibility, and biologically meaningful spatial organization relative to larger regions, while offering increased spatial specificity.

SLACS enabled reproducible analysis at this few-cell regime. At low input, successful tissue recovery becomes a major determinant of workflow performance because incomplete collection leads to loss of entire samples. Although UV-LCM yielded slightly higher protein identifications in successfully collected samples, SLACS provided substantially more consistent recovery of small tissue regions and reduced sample dropout. This improved collection robustness likely arises from the single-pulse release mechanism and the use of membrane-free metal oxide-coated slides, which reduce failure modes associated with perimeter cutting and polymer membrane detachment. At the same time, efforts to improve collection efficiency and reduce sample loss in UV-LCM workflows are also ongoing (Klingeberg *et al*., 2026) and will likely further increase the throughput and robustness of extraction-based spatial proteomic workflows. Together, these advances highlight the importance of continued optimization of tissue isolation workflows as spatial proteomics moves toward smaller and more spatially restricted tissue areas. SLACS should be viewed not only as an alternative isolation technology, but as an enabling strategy for robust operation at small sampling-unit size.

Our results indicate that biologically informative spatial proteomics can be achieved without necessarily operating at single-cell input. Although single-cell sampling provides maximal spatial specificity, cells differ widely in size, morphology, protein content, and contribution from local cellular processes. This is particularly relevant in complex tissues such as the cerebral cortex, where neurons, glial cells, axons, dendrites, and extracellular material contribute very different amounts of protein to the sampled area. For MS-based proteomics, which depends on the amount of analyzable material, a single cell therefore may not always constitute the most informative analytical sampling unit, particularly in heterogeneous tissues. In addition, normalization in single-cell proteomics is inherently challenging, as it is difficult to load equivalent amounts of protein from individual cells into the mass spectrometer, especially from different cell types, further complicating quantitative comparisons. Few-cell sampling units may provide a more practical compromise, preserving spatial information while improving robustness and proteome depth.

The mouse somatosensory cortex provided a stringent test of this concept. Cortical layers contain distinct cellular compositions and functional programs organized over fine spatial scales. SLACS- based sampling of approximately 60-cell regions generated deep, reproducible, layer-resolved proteomic profiles, recapitulating known cortical layer markers, superficial-layer synaptic enrichment, and deep-layer myelin-associated gradients. Importantly, these spatial proteome features were largely preserved when sampling-unit size was reduced to approximately 6 cells. This indicates that few-cell sampling captures biologically meaningful tissue organization rather than simply producing lower-depth proteomes.

At the same time, our data define the limitations of reducing sampling-unit size. Moving from approximately 60-cell to approximately 6-cell areas reduced proteome depth and preferentially affected detection of lower-abundance proteins. Thus, increasing spatial resolution comes at a measurable sensitivity cost. The key practical question is therefore not how small spatial proteomics can be pushed, but which sampling-unit size provides sufficient biological information with acceptable robustness and depth. In the systems tested here, few-cell sampling units offered a favorable balance between spatial specificity, proteome coverage, reproducibility, and collection success.

This work complements recent advances in extraction-based spatial proteomics, including DVP and cell type-resolved proteomic workflows (Mund *et al*., 2022; Rosenberger *et al*., 2023), which demonstrate the power of visually guided tissue isolation combined with ultra-sensitive MS. Our study focuses on the experimental trade-offs that emerge as sampling units become smaller in extraction-based spatial proteomics. As the field continues to improve MS sensitivity, sample preparation, and workflow automation, defining sampling strategies will become increasingly important for applications involving rare cell populations, spatially restricted microenvironments, and high-resolution tissue mapping.

In summary, our results illustrate how the balance between spatial specificity and proteome depth can be experimentally evaluated in extraction-based spatial proteomics. In the systems investigated here, few-cell sampling provides a practical compromise between spatial resolution, proteome depth and robustness, while SLACS enables reliable access to this sampling regime.

## Methods

### Animals

The animals used in this study were male C57BL/6J mice at ages P35 (liver) and P60-90 (brain), bred in-house at the animal facility of the Max Delbrück Center for Molecular Medicine. The animals were housed in standard Makrolon type II cages under a 12-hour light/dark cycle within a temperature-controlled environment (22 ± 2 °C) with 40-60 % humidity. Food and water were provided *ad libitum*. All experimental procedures described complied with institutional and European animal welfare guidelines and were approved by the Landesamt für Gesundheit und Soziales Berlin, Germany.

Animals were euthanized by an intraperitoneal administration of ketamine and xylazine (500 and 50 mg/kg, respectively), followed by sequential transcardiac perfusion with ice-cold phosphate-buffered saline buffer and 4% paraformaldehyde (PFA). Liver or brains were carefully dissected, post-fixed overnight in 4% PFA and subsequently cryoprotected in 30% sucrose (prepared in phosphate-buffered saline buffer) for 3 days. Liver tissue was then embedded in optimal cutting temperature medium (Tissue-Tek O.C.T., Sakura, 4583) and frozen on dry ice, while the brains were snap-frozen in methylbutane on dry ice. Tissues were then stored at -80 °C until further processing.

### Cryosectioning and tissue preparation

PPS-membrane (polyphenylene sulfide) FrameSlides (4.0 μm, MicroDissect, MDE5P80WK) and Cosmoslides (Meteor Biotech, Seoul, Republic of Korea; Cat. No. MET10102) were coated with Poly-L-Lysine (PLL, Sigma-Aldrich, P8920) for 5-10 min at room temperature, dried overnight and directly used for cryosectioning. The Cosmoslide consists of a glass support bearing a thin, optically transparent layer of metal oxide-coating that absorbs near-IR light; the composition and fabrication specifications of this layer are proprietary to the manufacturer. Brains were carefully mounted on sample holders and slightly covered with O.C.T. medium before sectioning. Liver tissues were mounted on sample holders within their embedded blocks. Tissue sections (5 μm (liver) or 10 μm (brain) thickness) were cut on a cryostat (CryoStar NX 70) and collected on the PLL-coated PPS-membrane FrameSlides (for UV-LCM) or on PLL-coated Cosmoslides (for SLACS). The tissues were stained with Hoechst (1:1000 in PBS, Thermo Fisher Scientific, 62249) and imaged on a Leica DMI6000 widefield fluorescence microscope. Tissues were stored at -20°C until further processing.

Throughout the manuscript ‘Cell-equivalent numbers’ refer to tissue areas estimated to contain the indicated number of cells based on tissue morphology and average cell density.

### LCM

Tissue areas were isolated with a LMD7 laser microdissection system (Leica) with a 20x dry objective in brightfield mode at 30°C with the following laser settings: power 60, aperture 17, speed 15, middle pulse count 1, final pulse 3, head current 51%, pulse frequency 1728 and offset 110. The isolated tissue areas were collected directly in low-bind 96-well plates and frozen at -20°C until further processing.

### SLACS

Tissue areas were isolated on a CosmoSort prototype instrument (Meteor Biotech, Seoul National University, Seoul, Republic of Korea), the direct predecessor of the commercial CosmoSort system (Cat. No. MET0001). The prototype supports isolation of regions with lateral dimensions between 1 μm and 1 mm, and all regions reported here were released with a single laser pulse. Hoechst-stained sections mounted on PLL-coated Cosmoslides were imaged on the instrument through a 20X objective, and regions of interest (ROI) were defined on the acquired images using the SLACS operating software. Each ROI was released from the slide by a single near-IR laser pulse (1,064 nm) delivered through the glass support. The released tissue areas were collected directly in low-bind 96-well plates and frozen at -20°C until further processing.

### Sample preparation for LC-MS

Sample plates were thawed at room temperature and the tissue regions were flushed to the bottom of the well with 10 μl acetonitrile (ACN), followed by evaporation with a speedVac concentrator for 10 minutes at 45°C. 3 μl of lysis buffer (0.1% DDM, 5 mM TCEP, 20 mM CAA, 100 mM TEAB) was added to all wells containing tissue regions. The plates were centrifuged at 2000 x g for 1 min and incubated at 95°C for 1 hour. Then, the plates were centrifuged again at 2000 x g for 1 min and 1 μl of Trypsin/LysC mix (4 ng, Promega, V507A) in 100 mM TEAB was added directly on top of the lysis buffer. After another centrifugation, the plates were incubated overnight at 37°C in a thermocycler. To stop the digestion, 0.1% formic acid (FA) was added to the samples.

Peptides were desalted and purified using Evotips (Evosep, EV2013). Evotips were first washed with 20 μl buffer B (0.1% FA in ACN), then soaked in isopropanol for 10 seconds and washed with 20 μl buffer A (0.1% FA in H2O). 10 μl of buffer A was added to the samples, pipette mixed and transferred to the evotips. Another 10 μl of buffer A was added to the sample wells, pipette mixed and transferred on top of the sample in buffer A from the previous step. The Evotips were centrifuged and washed twice with 20 μl of buffer A, then the peptides were eluted with 20 μl of buffer B. All centrifuge steps were performed at 800 x g for 1 min at room temperature. After elution, buffer B was evaporated through speedVac centrifugation at 45°C and samples were stored at - 20°C until LC-MS analysis.

### LC-MS analysis

Samples were thawed and resuspended in 4.2 μl loading buffer (0.1% FA in 3% ACN), of which 4 μl was injected. Samples were analyzed using a Vanquish Neo UHPLC (Thermo Fisher Scientific) coupled to an Orbitrap Astral MS (Thermo Fisher Scientific) operated in data-independent acquisition (DIA) mode. Peptides were separated on a 75 μm x 25 cm analytical column packed in-house with 1.9 μm C18 resin (ReproSil-Pur 120 C18-AQ, Dr Maisch).

Buffer A consisted of 0.1% FA in 3% ACN, and Buffer B consisted of 0.1% FA in 90% ACN. Peptides were separated using the following gradient of Buffer B: 2% to 7% within 1 min, then increased to 20% over 10 min and 30% over 7.5 min, followed by 60% over 2 min and 90 over 0.1 min. The column was washed for 4 min before re-equilibration. The total run time was 25 min.

The Orbitrap Astral MS was operated in positive ion mode with a spray voltage of 2.2 kV and a capillary temperature of 280°C. Full-scan MS spectra were acquired at a resolution of 240,000 over a mass range of 380-1100 m/z, with a maximum injection time of 100 ms. DIA spectra were acquired in the Astral mass analyzer with 8 m/z windows covering 380-980 m/z, a normalized AGC target of 500%, a normalized collision energy of 25, and a maximum injection time of 14 ms.

One mouse liver experiment was analyzed using an EASYnLC-1200 (Thermo Fisher Scientific) coupled to a TimsTOF SCP (Bruker) operated in DIA mode. Here, peptides were separated using a linear gradient from 7-30% Buffer B (0.1% FA in 90% ACN) over 14 min, increased to 60% over 1 min, washed at 90% buffer B for 1.5 min, then decreased to 50% over 3.5 min. The total run time was 21 min.

For dia-PASEF acquisition, the mass spectrometer was operated in high sensitivity mode with the following parameters: 8 dia-PASEF scans, each with 3 ion mobility windows covering 400-1000 m/z range with 25 Th windows, an ion mobility range of 0.64 to 1.37 Vs/cm^2^, an accumulation and ramp time at 100 ms, the capillary voltage set to 1600 V and the collision energy was ramped linearly as a function of ion mobility from 20 eV at 1/K0 = 0.6 Vs/cm^2^ to 59 eV at 1/K0 = 1.6 Vs/cm^2^.

### Data-analysis

Protein identification was performed with DIA-NN (v2.2.0, Demichev *et al*., 2019) using the mouse reference proteome (UniProt UP000000589) as the database and trypsin specificity, allowing up to two missed cleavages and three variable modifications per peptide. Peptides of 7-52 amino acids with precursor charges 2-4 were considered. Fragment ions from 100-1800 m/z were analyzed with mass accuracies of 10 ppm (Astral) or 15 ppm (TimsTOF) and a scan window of 6.

Match-between-runs and cross-run normalization were disabled. Downstream data analysis was performed in Python (v3.12.8) using Jupyter Notebook (v7.3.2).

### Mouse liver samples

To calculate collection efficiencies (**Fig 1f**), mouse liver samples with a sampling-unit size of 6,000 μm^2^ (equivalent to approximately 6 cells) with fewer than 1,000 identified protein groups were classified as failed collections. For protein overlap analyses (**Fig 1h**), proteins were considered detected if identified in at least 70% of replicates within a condition. Protein abundance comparisons (**Fig 1i**) were performed using median log_10_-transformed protein intensities across replicates for all shared protein groups, irrespective of detection frequency. Similarity between the 100-cell and 6-cell datasets was assessed using Pearson correlation of protein abundances. Sample-to-sample correlations (**Extended Data Fig 1e**) were calculated using pairwise Pearson correlation coefficients based on log_10_-transformed protein intensities, with zero intensity values treated as missing prior to transformation.

### Mouse brain samples

One low-performing sample in the 6-cell mouse brain dataset was excluded because fewer than 1,000 protein groups were identified, resulting in five replicates for one cortical layer (L6) and six for all remaining conditions. Protein groups with more than one missing value in every cortical layer were removed, retaining only those with sufficient quantified values in at least one condition.

Coefficients of variation (CVs) were calculated (**Extended Data Fig 2i**) from normalized, non-log-transformed protein intensities across biological replicates within each cortical layer. Zero intensity values were treated as missing values before calculation. Protein abundance comparisons (**Extended Data Fig 2g**) were performed using median log_10_-transformed protein intensities across replicates for all shared protein groups, irrespective of detection frequency. Similarity between the 60-cell and 6-cell datasets was assessed using Pearson correlation of protein abundances.

For subsequent analyses, protein intensities were log_2_-transformed. Missing values were imputed separately within each cortical layer using a Gaussian missing-not-at-random (MNAR) imputation approach. Imputed values were randomly sampled from a normal distribution centered at the 0.5^th^ percentile of the observed log_2_-transformed intensities within each cortical layer, with a standard deviation equal to 0.3 times the global standard deviation of the observed intensities. Data were median normalized across samples to correct for systemic differences in protein abundance. PCA (**Fig 2c and 2d; Extended Data Fig 2c**) was performed on the log2-transformed, median-normalized and imputed protein intensities. Before PCA, each protein group was standardized across samples to zero mean and unit variance.

Differential protein abundance analysis was performed using the limma package (v3.62.2) in R (v4.4.2) via an R-Python interface. Linear models were fitted to normalized protein intensities followed by empirical Bayes moderation. Six pairwise comparisons were performed between cortical layers (L2/3 versus L4, L2/3 versus L5, L2/3 versus L6, L4 versus L5, L4 versus L6, and L5 versus L6). *P* values were adjusted using the Benjamini-Hochberg false discovery rate (FDR). Proteins were considered significantly differentially abundant at an adjusted *P* value < 0.05 and absolute log₂ fold change > 0.58. Heatmaps were generated using the protein groups selected for each analysis. For visualization, the abundance values of each protein group were standardized by z-scoring across all samples.

Protein overlap analyses (**Extended data Fig 2d and 2e**) were performed using protein groups identified in each dataset after quality control and filtering. For differential abundance overlap analyses, proteins showing significant differential abundance in at least one pairwise comparison between cortical layers (adjusted *P* < 0.05 and absolute log₂ fold change > 0.58) were compared between datasets.

Proteins showing significant differential abundance in at least one pairwise comparison across cortical layers were clustered using k-means clustering implemented in scikit-learn (Python). Protein abundance values were row-wise z-score standardized across samples prior to clustering. The number of clusters (k = 5) was selected based on elbow and silhouette score criteria. Clustering was performed using the Kmeans algorithm with a fixed random state (random_state = 42) to ensure reproducibility. Cluster centroids and member profiles were used for downstream visualization and functional enrichment analysis.

GO-term enrichment analysis was performed in Cytoscape (v3.10.3) using CleuGO (v2.5.10) with Biological Process ontologies and GO-term fusion enabled. Enrichment analyses were performed using proteins showing significant differential abundance in at least one pairwise comparison between cortical layers, proteins shared or unique between datasets, or proteins assigned to individual k-means clusters, as indicated for each figure. Only terms with a Bonferroni corrected *P* ≤ 0.05 were considered, using all DIA-NN identified protein groups as the reference background. Corrected *P* values were -log_10_-transformed for visualization.

## Supporting information

Meijer_Supplementary Table 1

Meijer_Supplementary Table 2

## Acknowledgments

We thank Fabian Coscia and Di Qin for helpful scientific discussions and for their advice and technical guidance on the LCM workflow. We also thank all group members of the Piazza laboratory, the proteomics community and the advanced light microscopy technology platform at MDC Berlin for helpful scientific exchange and technical support. This work was supported by the Initiative and Networking Fund of Helmholtz Young Investigators program of the Helmholtz Association (funding code: VH-NG-1535), by the Deutsche Forschungsgemeinschaft (DFG, grant agreement No 535634027) and by the European Research Council (ERC) under the European Union’s Horizon 2020 research and innovation program (grant agreement ERC-STG No 948544) to IP. MM acknowledges support from the Alexander von Humboldt foundation and from Svenska Sällskapet för Medicinsk Forskning (SSMF; PG-23-0364-H-01). IP also acknowledges funding from the Wallenberg Academy Fellow program (KAW 2023.0053), Stockholm University and Karolinska Institutet. VP acknowledges funding from an extension Wallenberg Academy Fellowship (KAW 2021.0167), the Swedish Research Council (VR 2020-01480 and 2024-03210) and Karolinska Institutet (SciLifeLab Fellowship, SFO, and KI funds). Work at Meteor Biotech was further supported by the National Research Foundation of Korea (NRF), funded by the Korean government (MSIT; grant no. RS-2024-00454407 to A.L.); by the Technology Innovation Program and the Industrial Strategic Technology Development Program of the Ministry of Trade, Industry and Energy (MOTIE), Republic of Korea (grant nos. 20024391, RS-2024-00451981 and RS-2024- 00508420 to A.L.); and by the Korea–US Collaborative Research Fund (KUCRF), jointly funded by the Ministry of Science and ICT and the Ministry of Health & Welfare, Republic of Korea (grant no. RS-2024-00468338 to A.L.).

---

**Extended Data Figure 1.**
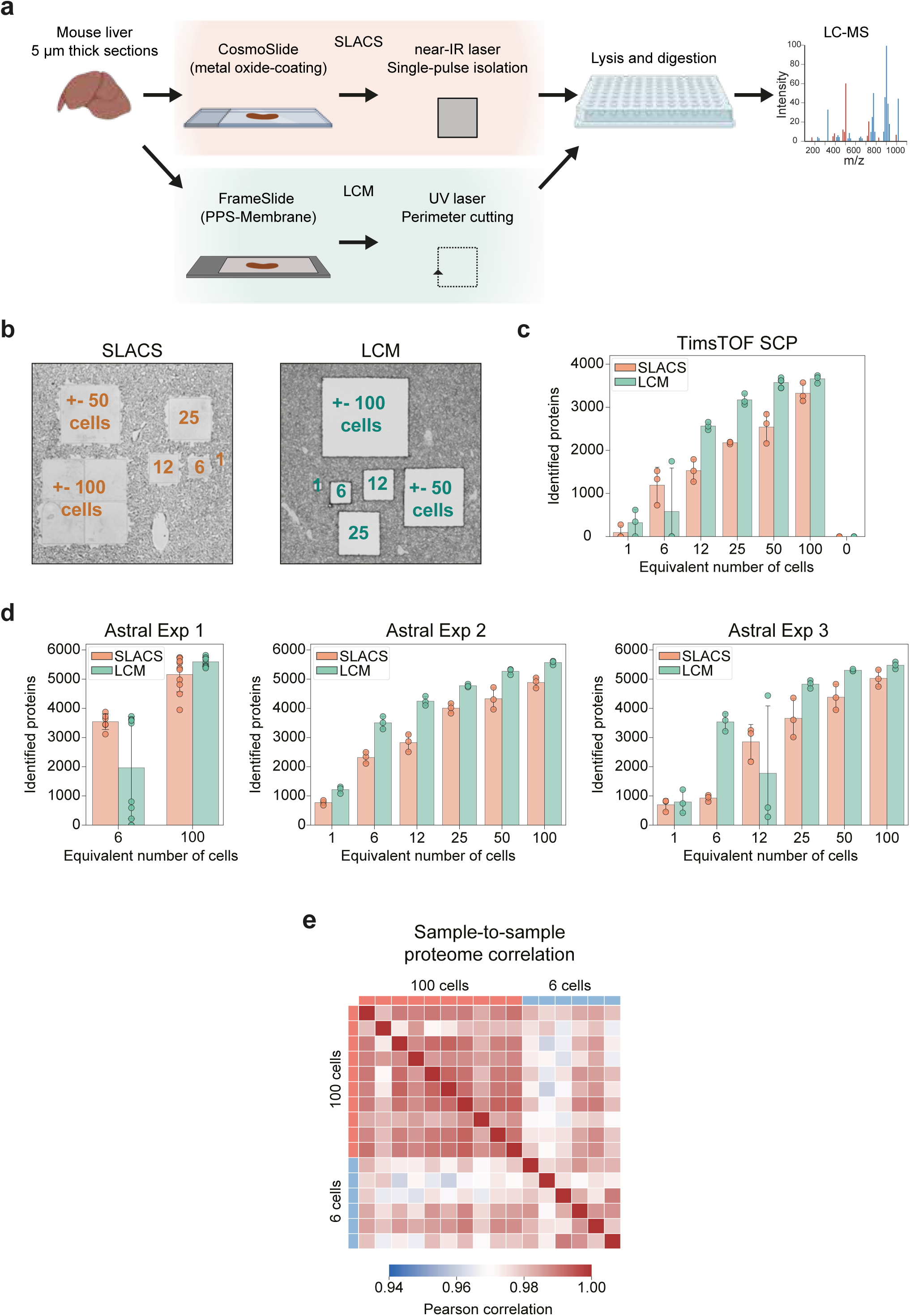
**a**, Workflow for isolation of defined tissue areas from mouse liver using SLACS or UV-LCM. In UV- LCM, tissue areas are isolated by perimeter cutting from polymer membrane slides, whereas SLACS uses single-pulse near-IR laser activation to release tissue areas from metal oxide-coated slides. Isolated areas are collected into 96-well plates for in-well lysis and digestion prior to DIA-MS analysis. **b**, SLACS and UV-LCM isolation of liver tissue areas of 600 μm^2^, 6,000 μm^2^, 12,500 μm^2^, 25,000 μm^2^, 50,000 μm^2^ and 100,000 μm^2^, equivalent to approximately 1, 6, 12, 25, 50, and 100 cells. Wide-field image of the liver tissue after isolation. **c**, Number of identified protein groups per collected input size using SLACS (orange) or UV-LCM (green), measured on a Bruker TimsTOF SCP mass spectrometer. Bars represent the mean and error bars indicate SD (n=3). **d**, Number of identified protein groups per collected input size using SLACS (orange) or UV-LCM (green), measured on an Thermo Fisher Orbitrap Astral mass spectrometer. Data represent 6-10 replicates (Exp1) and 3 replicates (Exp2 and 3) per collected input size. Data are from three experiments processed and measured on different days. Bars represent the mean and error bars indicate SD. **e**, Heatmap of pairwise Pearson correlation coefficients between individual SLACS samples collected at the 6-cell and 100-cell sampling-unit sizes. Correlations were calculated directly from log_10_-transformed protein intensities. Zero intensity values were treated as missing, and each pairwise correlation was calculated using proteins quantified in both samples. Data represent 6 replicates for the 6-cell sampling unit and 10 replicates for the 100-cell sampling unit.

**Extended Data Figure 2.**
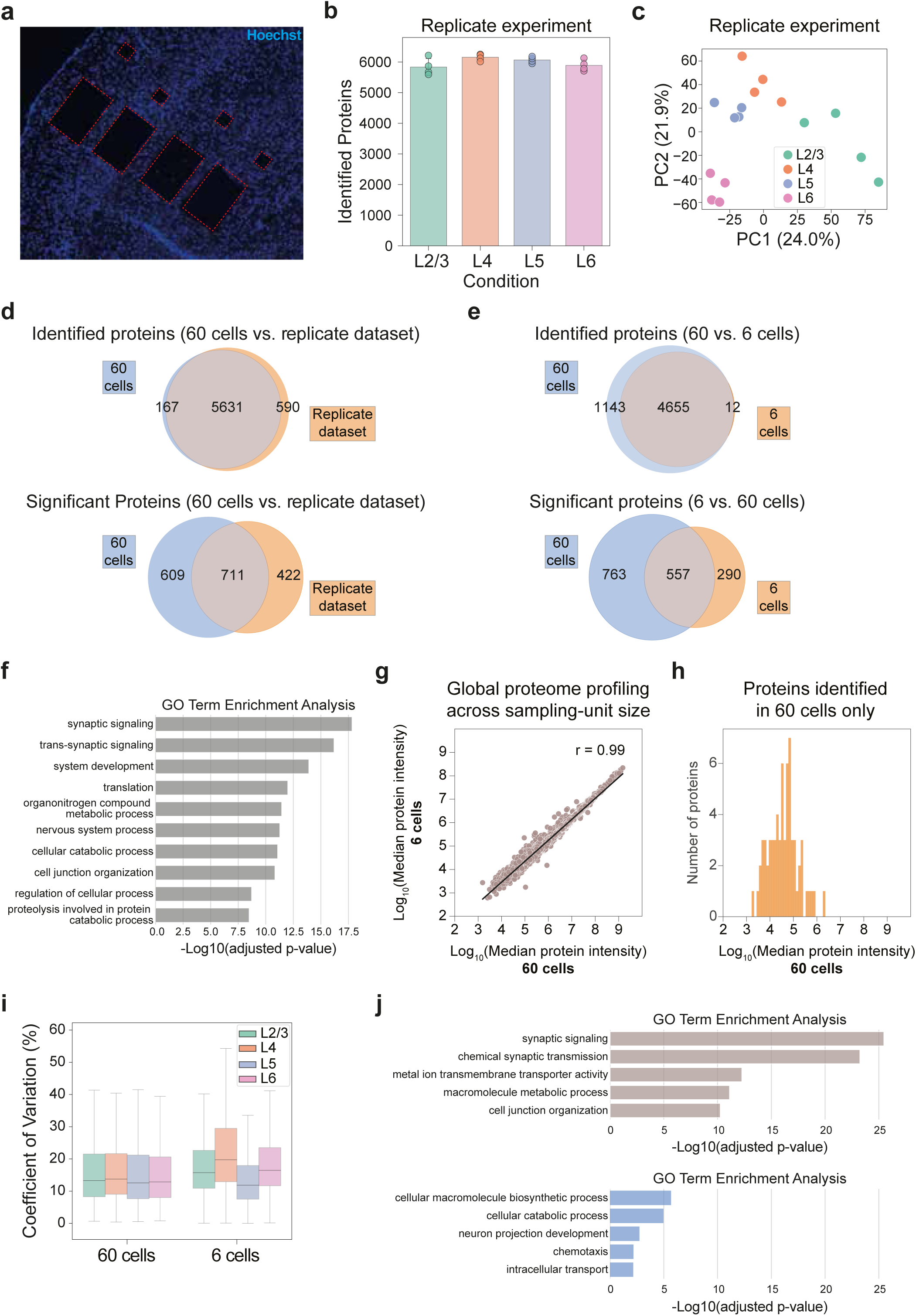
**a**, Widefield image of the somatosensory cortex following isolation of 60- and 6-cell areas. Hoechst staining is shown in blue. **b,** Number of identified protein groups per cortical layer in an independent replicate experiment (60-cell areas). Data represent 4 replicates per cortical layer. Bars represent the mean and error bars indicate SD. **c**, Principal component analysis of scaled protein abundances across cortical-layer samples at the 60-cell sampling unit size of the replicate experiment. Each point represents one replicate. **d**, Overlap of proteins identified in the 60-cell dataset and the replicate dataset for total proteins (top) and proteins showing significant differential abundance in at least one pairwise comparison between cortical layers, identified using linear models with empirical Bayes moderation (adjusted p < 0.05, Benjamini-Hochberg FDR correction, |log_2_ fold change| > 0.58) (bottom). Experiments are independent. **e**, Overlap of proteins identified in the 60- and 6-cell datasets for total protein groups (top) and proteins showing significant differential abundance in at least one pairwise comparison between cortical layers, identified using linear models with empirical Bayes moderation (adjusted p < 0.05, Benjamini-Hochberg FDR correction, |log_2_ fold change| > 0.58) (bottom). **f**, GO-term enrichment analysis of proteins showing significant differential abundance in at least one comparison between cortical layers in the 60-cell dataset. The top 10 GO terms are shown, ranked by significance (-log_10_ Bonferroni corrected *P* value). **g**, Scatter plot comparing median protein intensities measured with DIA-MS between 60- and 6- cell samples. Each point represents one protein quantified at both sampling-unit sizes using the median log_10_-transformed intensity across 5-6 replicates. Protein abundances were highly correlated across sampling-unit sizes (Correlation coefficient r = 0.99; Pearson correlation). **h,** Distribution of protein abundances for proteins identified in 60-cell samples but not in 6-cell samples. Intensities correspond to the median protein abundance in the 60-cell dataset. Proteins absent at the 6-cell sampling-unit size were predominantly of low abundance, indicating that reducing sampling-unit size primarily affects detection sensitivity. **i**, Protein-level coefficients of variation (CVs) across replicates for each cortical layer at the 60-cell and 6-cell sampling-unit sizes. CVs were calculated from normalized, non-log-transformed protein intensities. Center lines indicate the median, boxes the interquartile range, and whiskers extend to 1.5x the interquartile range. Outliers are not shown for clarity. **j**, GO-term enrichment analysis of proteins showing significant differential abundance in at least one comparison between cortical layers in both the 60- and 6-cell datasets (top) or only in the 60- cell dataset (bottom). The top five GO terms are shown, ranked by significance (-log_10_ Bonferroni corrected *P* value).

**Extended Data Figure 3.**
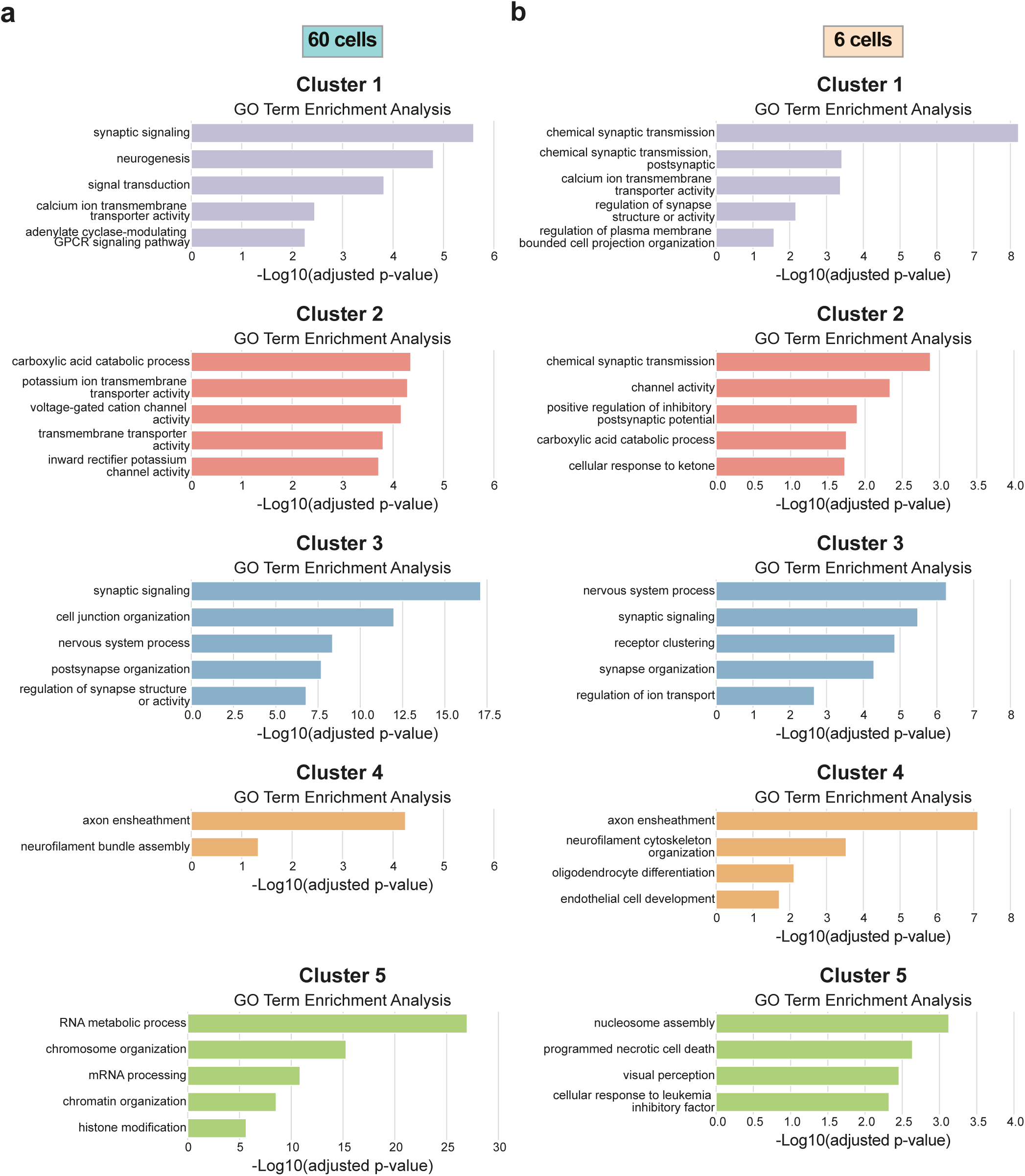
**a**, GO-term enrichment analysis of proteins in each cluster shown in Fig. 3a (60-cell dataset). The top five GO terms are shown, ranked by significance (-log_10_ Bonferroni corrected *P* value). **b**, GO-term enrichment analysis of proteins in each cluster shown in Fig. 3b (6-cell dataset). The top five GO terms are shown, ranked by significance (-log_10_ Bonferroni corrected *P* value).

Supplementary Table 1. Proteins showing significant differential abundance across cortical layers in the 60-cell dataset.

Supplementary Table 2. Proteins showing significant differential abundance across cortical layers in the 6-cell dataset.

